# Mechanical tuning of replication stress tolerance and genomic stability through the checkpoint mediator Mrc1

**DOI:** 10.64898/2026.05.14.725212

**Authors:** Adam J. Timmerman, Rowan E. Brady, Via M. Lawson, Joseph A. Stewart, Juan Lucas Argueso, Grant D. Schauer

## Abstract

Mrc1 is a replication fork component that mediates communication between the replication checkpoint and fork progression. Here, we find that Mrc1 forms a dynamic mechanical linkage between CMG helicase and DNA Polymerase ε that balances fork stability and flexibility. In its coupled state, Mrc1 bridges CMG helicase and the Pol2 C-terminal domain of Polymerase ε to maintain helicase–polymerase coordination, promoting tolerance to replication stress but constraining fork remodeling. Upon checkpoint activation, Mec1/Rad53-dependent phosphorylation of Mrc1 shifts this equilibrium toward a loosened state, reducing coupling and enabling fork flexibility and lower mutation rate at the cost of reduced stress resistance.

## INTRODUCTION

Genetic material within the cells of every living organism must be duplicated in a manner that minimizes accumulation of mutations while also proceeding expeditiously (Yates et al. 2025). During eukaryotic replication, CMG helicase (<u>C</u>dc45-<u>M</u>cm2-7-<u>G</u>INS) encircles the leading strand template and is tightly coupled to DNA polymerase (Pol) ε. The CMG-Pol ε linkage has been shown to be necessary for both replication initiation and elongation (Xu et al. 2023; Handa et al. 2012). Continuous monitoring of the replisome ensuring enzymatic coupling between helicase and polymerase is therefore critical for survival.

During S-phase, genome integrity is maintained by an ensemble of factors which constitute the essential replication checkpoint (Yates et al. 2025). This checkpoint can be activated by a variety of replication stressors like lesions or gaps present on the template strand or by limited pools of deoxyribonucleotides, to name a few (Zeman and Cimprich 2014). An oft-used means of uncoupling forks employs hydroxyurea (HU) as a stressor, which destabilizes replicative Pols and depletes dNTP levels (Shaw et al. 2024). In all forms of stress, the replication fork becomes uncoupled when CMG helicase and Pol ε separate, triggering checkpoint activation, which in turn protects replication forks from collapse while replication stress is mitigated. In *Saccharomyces cerevisiae*, checkpoint activation occurs upon recruitment of Mec1 kinase and its regulatory partner Ddc2 to extended tracts of RPA-bound single-stranded DNA resulting from uncoupling; the polymerase stalls, but the helicase continues to unwind the double-stranded DNA ahead of the fork (Rouse and Jackson 2002; Zou and Elledge 2003). Following its recruitment to ssDNA tracts, Mec1 activates the effector kinase Rad53 which, in addition to numerous regulatory effects, stabilizes stalled replication forks (Paulovich and Hartwell 1995; Lopes et al. 2001; Tercero and Diffley 2001).

Early studies of the S phase checkpoint determined that Pol2, the large polymerase subunit of the Pol ε holoenzyme, is necessary for Rad53 phosphorylation and checkpoint activation (Navas et al. 1995, 1996). Pol2 has an N-terminal domain (NTD) harboring polymerization activity and a catalytically inactive C-terminal domain (CTD) that nonetheless largely retains a polymerase fold (Goswami et al. 2018). Surprisingly, studies have shown that while deletions of Pol2^NTD^ can be moderately tolerated, deletion of Pol2^CTD^ is strictly lethal (Dua et al. 1999; Kesti et al. 1999). Pol2^CTD^ was later found to structurally scaffold Pol ε binding to CMG (Georgescu et al. 2017). Thus, while the role of Pol2 in the replication checkpoint response is established, the mechanism that couples it to activation is less clear.

Following findings regarding the role of Pol2 in the checkpoint, Mrc1 (Mediator of the Replication Checkpoint) emerged as a mediator of Rad53 activation by Mec1 through a mechanism that remains unclear. Initial studies identified Mrc1 as a member of the fork protection complex, along with Tof1 and Csm3 (Alcasabas et al. 2001; Katou et al. 2003; Naylor et al. 2009; Bando et al. 2009). The presence of Mrc1 is required for normal progression through S-phase (Osborn and Elledge 2003; Hodgson et al. 2007) and, when added to *in vitro* replication reactions with pure proteins, increases replication rates to those observed physiologically by boosting fork speed about 1.5-fold (Osborn and Elledge 2003; Hodgson et al. 2007; Yeeles et al. 2017; Lewis et al. 2017, 2020). Scattered throughout Mrc1 (largely in the N-terminal half; **Fig. 2A**) are 17 serine or threonine residues followed by a glutamine residue (SQ/TQ), clusters of which are the canonical targets of Mec1 kinase activity (Traven and Heierhorst 2005). Pioneering studies of Mrc1 introduced the *mrc1*^*AQ*^ allele, in which the serine or threonine of all 17 SQ/TQ sites are mutated to alanine to prevent their phosphorylation, disabling the checkpoint-activating function of Mrc1 and increasing the sensitivity of cells to replication stressor drugs like HU and methyl methanesulfonate (Osborn and Elledge 2003). Interestingly, unperturbed *mrc1*^*AQ*^ cells progress through S-phase at the same rate as wild-type cells, demonstrating that the replication stimulation and S-phase checkpoint activation functions of Mrc1 are distinct and separable (Osborn and Elledge 2003).

Biochemical and genetic studies have provided critical glimpses into the mechanisms underlying the function(s) of Mrc1. In reconstituted kinase assays, Mrc1 was found to promote Mec1-Ddc2 association to Rad53, enhancing kinase activity (Chen and Zhou 2009). Truncation mutants of the *mrc1* allele have also been used to map the domains required for replication stress resistance, and, by extension, checkpoint competence (Naylor et al. 2009; Chen and Zhou 2009). Studies have shown that Mrc1 interacts with several components of the eukaryotic replisome, notably CMG helicase and Pol ε (Lou et al. 2008, Zhang et al. 2018), requiring Tof1 and Csm3 to recruit Mrc1 to CMG and to stimulate replication speeds (Yeeles et al. 2017; McClure and Diffley 2021). Moreover, the nature of Mrc1 stimulation of replication fork speed is an area of active study, with several groups demonstrating that Mrc1 enhances CMG unwinding rates (Gan et al. 2017; Serra-Cardona et al. 2021; Devbhandari and Remus 2020; McClure and Diffley 2021; He and Zhang 2022), and at least one group demonstrating that Mrc1 can stimulate Pol ε activity directly (Zhang et al. 2018). Additionally, one study indicates that phosphorylation of Mrc1 by Rad53 is necessary and sufficient to attenuate CMG unwinding rate (McClure and Diffley 2021) while another suggests that phosphorylation of CMG by Rad53 is sufficient to reduce fork speeds (Devbhandari and Remus 2020). It has also been postulated that in regulating CMG unwinding rate, Rad53 plays a role in coupling unwinding and synthesis activity between leading and lagging strands (Gan et al. 2017; Serra-Cardona et al. 2021). Despite these advances, the highly charged and disordered nature of Mrc1 presents challenges to its structural determination alone or in complex with other replisome components. Significant effort has been made to resolve its structure, but determination has remained elusive with only a handful of interaction sites identified by cross-linking mass spectrometry (XL-MS) that localize regions of Mrc1 binding to CMG (Baretić et al. 2020). That study, which determined the structure of CMG with Tof1, Csm3, and Ctf4, approximated Mrc1 orientation along the DNA-parallel axis of the replisome by identifying crosslinks of the Mrc1 N-ter region to Tof1, situated at the helm of CMG that unwinds the dsDNA duplex, and of the Mrc1 C-ter region to Ctf4, located at the trailing end of the fork where Pols ε and α are generally localized. More recently, it was shown in *Schizosaccharomyces pombe* that Mrc1 has yet a third role as a histone chaperone, with this activity residing in the N-terminal region (Yu et al. 2024). Still, there is a substantial gap in understanding how Mrc1 enhances replication rates and facilitates activation of the replication checkpoint, as well as how these two functions might be related to each other. Taking this context in sum and given that Mrc1 was shown by co-immunoprecipitation and yeast-two-hybrid to bind CMG and Pol ε separately (Lou et al. 2008), we asked whether Mrc1 can indeed couple the activity of these principal fork components.

In this report, we use *in vitro* biochemical and single-molecule reconstitution of replication fork uncoupling to demonstrate that Mrc1 mechanically tethers CMG helicase and Polymerase ε during replication. Using multiparallel predictive modeling, we predict and validate an interaction site between the C-terminal region of Mrc1 and the noncatalytic Pol2^CTD^ involving a conserved salt bridge, providing a structural framework for the concerted actions of Mrc1 on the replication fork machinery. We show using quantitative recombination and point mutation assays that *mrc1* mutations which shift the equilibrium towards helicase-polymerase uncoupling improve the overall baseline genomic stability of budding yeast, while conversely increasing sensitivity to stress. Although genomic stability is enhanced overall by these mutations, forced uncoupling differentially affects distinct classes of genome instability outcomes, suggesting a redistribution between replication-associated error events and repair outcomes following fork perturbation. We relate these findings to varied degrees of activation of the S-phase checkpoint determined by Rad53 phosphorylation. Our results shed light on the role of Mrc1 in fine-tuning the checkpoint response to balance replisome plasticity and stress tolerance with genomic stability and replication efficiency.

## RESULTS AND DISCUSSION

### Mrc1 physically, functionally, and dynamically couples CMG and Pol ε

We first asked whether Mrc1 couples CMG and Pol ε during replication elongation. We developed “replication uncoupling” assays by modifying our *in vitro* primer extension assays, which use numerous purified proteins to reconstitute the minimal leading strand replisome. The leading strand is enzymatically decoupled from lagging strand machinery and can thus be studied in isolation by following the fate of a single leading strand primer (Schauer et al. 2017; Schauer and O’Donnell 2017). In these reactions, we first loaded CMG onto a forked DNA duplex (a 2.8 kb duplex ligated to a noncomplementary fork) in the presence or absence of the Mrc1 in the form of the Mrc1-Tof1-Csm3 (MTC) complex, followed by Pol ε, RFC clamp loader, and PCNA clamp. We initiated fork uncoupling by adding the ATP required for CMG unwinding along with a limited set of dNTPs (dATP and dCTP) and RPA—during this phase the polymerase abortively attempts to incorporate miscognate nucleotides and remains in the polymerase-competent enzymatic mode (as opposed to the proofreading mode) at the primer 3’-terminus while CMG continues to progressively unwind the duplex. After a designated uncoupling time, Δt, we then added the remaining requisite dNTPs (dTTP and dGTP). Since the duplex will reanneal in the wake of CMG and since Pol ε lacks significant strand displacement activity, we predicted that the fork would become progressively uncoupled with increasing Δt (see schematic in **Fig. 1A**). Indeed, uncoupling reactions were severely hindered compared to normal replication reactions, especially without Mrc1 present, resulting in largely stalled extension products (e.g., compare **Fig. 1B** to **Fig. S1A**). In the presence of WT Mrc1, a modest amount of coupling could be seen, leading to a visible full-length product (see **Fig. S1B** for enhanced contrast). However, adding Mrc1^AQ^ resulted in a strong and predominately full-length band (**Fig. 1B**, quantified in **Fig. 1C)**. Since the AQ mutation did not alter the fork stimulating activity of Mrc1 (**Fig S1A)**, we attributed this effect to more efficient CMG-Pol ε coupling (**Fig 1A**). To determine whether this coupling was physical, we next performed nearly identical experiments at the single-molecule level except for the use of a shorter biotinylated template and withholding the final dNTPs to avoid replication completion and replisome runoff. We designed a Fluorescence Resonance Energy Transfer (FRET) uncoupling assay using CMG labeled at the C-ter of Cdc45 (CMG^555^) and Pol ε labeled at the C-ter of Pol2 (Pol ε^655^) predicted to have high FRET when coupled (**Fig. 1D, Fig. S2A**). Site-specific dye labeling did not affect activity of CMG or Pol ε (**Fig. S2B**). Monitoring FRET between CMG^555^ and Pol ε^655^, we observed low FRET without Mrc1, a modest increase with WT Mrc1, and high FRET with Mrc1^AQ^, matching our biochemical observations (**Fig. 1D)**, also in agreement with previous Y2H evidence that Mrc1^AQ^ binds Pol2^CTD^ more tightly (Lou et al. 2008). Generally, coupling FRET trajectories appeared to be more dynamic either without Mrc1 or with WT Mrc1 than in the presence of Mrc1^AQ^ (**Fig. S2C**). Since Mrc1^AQ^ consistently coupled replication better than WT Mrc1, the results are consistent with our WT Mrc1 carrying a basal amount of phosphorylation, which was supported by tandem mass spectrometry on purified Mrc1 (Supplementary Info). Since Mrc1^AQ^ lacks the capacity for Mec1-dependent phosphorylation at SQ/TQ residues, we interpret this to mean that Mrc1—and by corollary, fork coupling—can be fine-tuned by phosphorylation status (**Fig 1D** schematic), in support of a previous prediction (Lou et al. 2008).

**Figure 1.**
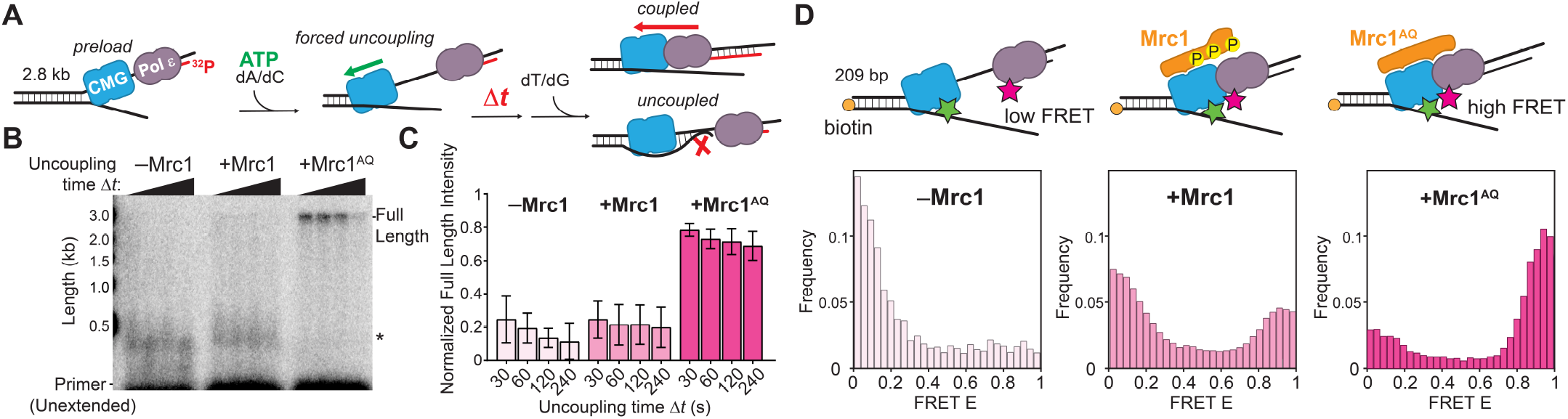
Mrc1 mechanically tethers Pol ε to CMG and is tunable by phosphorylation status. **A)** Forced uncoupling assay schematic. CMG and Pol ε are loaded at the fork, including Mrc1 where indicated. Tof1, Csm3, RPA, RFC, and PCNA were present in all reactions but are omitted for clarity. Uncoupling assay staging is described in Results and Discussion. **B)** Example autoradiograph of primer extension products from forced uncoupling assay resolved by alkaline agarose gel. Full length (2.8 kb) product indicates a coupled leading strand replisome. The asterisk indicates stalled regions of Pol ε. Results from 3 experiments were quantified in **C)** showing the mean ±STD of the full-length intensity normalized to all other extended products. Uncoupling times are indicated. Results were statistically significant between corresponding timepoints of +Mrc1^AQ^ and either –Mrc1 or +Mrc1 conditions and were n.s. between any other conditions according to a two-tailed t-test between values. **D)** Single-molecule FRET uncoupling assay. Top: schematic. FRET occurs when CMG bearing LD555 (green star) is close to Pol ε labeled with LD655 at the Pol2 CTER (red star). The reaction is staged like the biochemical uncoupling assay except the second round of cognate dNTPs is not added. Bottom: FRET histograms are shown for post-uncoupling reactions for –Mrc1 (n=417), +Mrc1 (n=841), or +Mrc1^AQ^ (n=420) conditions.

### The C-terminal region of Mrc1 differentially controls stress tolerance and mutation propensity

Intrigued by observing Mrc1-directed replisome coupling, we initially tried to map Mrc1 binding to Pol ε via XL-MS but did not identify any crosslinks (not shown). Recently, however, predictive modeling of *S. pombe* Mrc1 identified a putative Pol2 interaction domain, though nothing of this interaction was revealed beyond its identification (Yu et al. 2024). Using multiple sequence alignment (MSA; see Methods), this region (880-945) corresponds to residues 926–1001 in *S. cerevisiae*. We therefore wanted to test the phenotype of cells lacking the region known to be important for checkpoint activation and for interacting with CMG (Naylor et al. 2009; Baretić et al. 2020) (*mrc1*^1–720^) vs. cells lacking a region in Mrc1 we reasoned might more specifically bind Pol ε (*mrc1*^1–946^; see **Fig. 2A** for map of these regions). Consistent with previous results showing impaired checkpoint in cells with *mrc1* truncated before residue 820 (Naylor et al. 2009), *mrc1*^1–720^ cells displayed notably increased HU sensitivity. Surprisingly, however, *mrc1*^1–946^ cells exhibited increased HU tolerance compared to WT in our hands (**Fig. 2B**). Adding CEN (single-copy) plasmids expressing the missing C-terminal domains, we saw that *mrc1*^721–end^ was able to modestly rescue WT and *mrc1*^1–720^ cells but did not further enhance the HU tolerance seen in *mrc1*^1-946^ cells. Expression of *mrc1*^947-end^ had no effect on cells lacking either CTER domain; however, it did enhance HU tolerance in WT cells (**Fig. 2B**). We did not find notably different expression or phosphorylation shifts of Rad53 that might explain these findings in a signaling context. Thus, we reasoned that at least part of Mrc1^947–end^ may be responsible for binding Pol2. Interestingly, contrary to single-copy expression, overexpression of either *mrc1* CTER domain appears harmful to cells and sensitizes them to HU, but only in a *mrc1*Δ background, which we interpret as unproductive coupling by this region in the absence of CMG stimulatory and/or checkpoint activity (**Fig. S4**).

**Figure 2.**
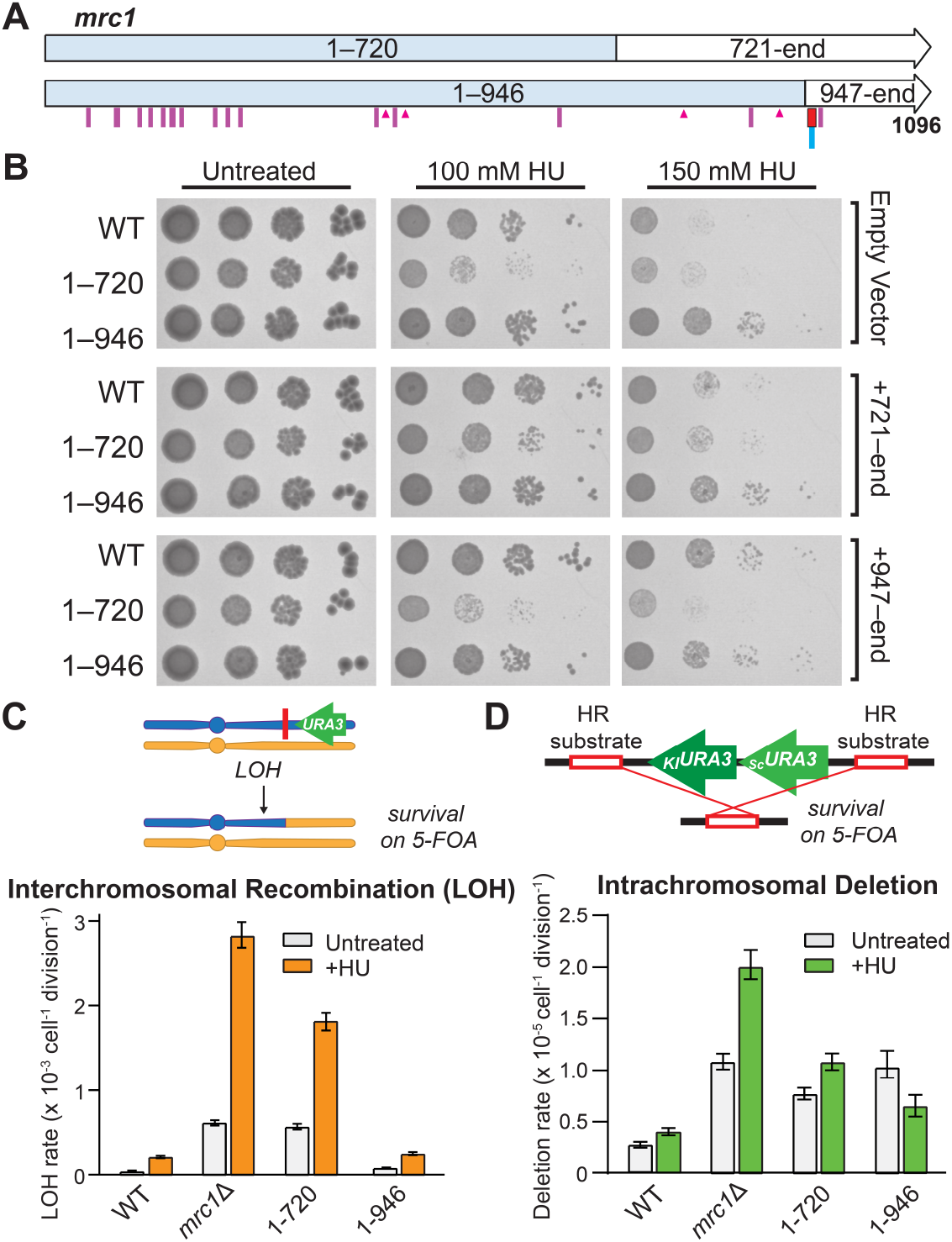
The C-terminal region of Mrc1 differentially controls stress tolerance and mutation propensity. **A)** Primary structure of Mrc1 highlighting the fragments discussed throughout this work. Purple lines indicate locations of SQ/TQ residues; triangles indicate CMG interaction residues previously identified by XL-MS (discussed in Introduction); red box and blue line respectively indicate N16 and K954, discussed in Fig. 3. **B)** Survival spot assays of the indicated MRC1 truncation mutants are shown in response 150 mM HU stress where indicated. Where indicated, cells were transformed with single-copy CEN plasmids expressing the specified Mrc1 fragment. **C)** LOH on Chr. 4 rates and **D)** intrachromosomal deletion rates on Chr. 4 for the indicated *MRC1* genotypes, in the presence or absence of 25 mM HU as indicated. Bars represent the maximum likelihood mutation rate ± 95% CI; significance was inferred where CIs do not overlap, corresponding to *p* < 0.05. The full mutation rate dataset is provided in Fig. S9.

Fork protection prevents collapse and the associated DNA break lesions that trigger recombination and genomic rearrangements during replication, especially under stress. Thus, we next asked whether lack of these Mrc1 regions could alter loss-of-heterozygosity (LOH) or deletion rates (see schematic; **Fig. 2C and Fig. 2D** respectively). Though we saw drastically increased LOH rates in both *mrc1*Δ and *mrc1*^*1-720*^ cells in the presence or absence of HU (12- and 8-fold without, and 13- and 12-fold with HU, respectively), the spontaneous LOH rate in *mrc1*^*1-946*^ cells was only moderately higher than WT without HU (1.3-fold), and was surprisingly unchanged in the presence of HU (**Fig. 2C**; see **Fig. S9A** for log-scale plot and full dataset). When we measured intrachromosomal deletion rates (**Fig. 2D**), we saw that HU treatment caused moderately higher rates compared to baseline in all *mrc1* backgrounds except *mrc1*^1–946^, where it was significantly lower than in untreated cells (**Fig. 2D**). We hypothesized that physical interaction with Pol2 may underlie the ability of this Mrc1 region to differentially control stress tolerance and mutation propensity during replication.

### Identifying a conserved interaction between Mrc1 and Pol2^CTD^

We wanted to know how Mrc1^947–end^ might interact with Pol2. We first attempted to structurally predict Mrc1 interactions with CMG and/or Pol2 with Alphafold 2 (AF2), but predictions were generally low confidence, disordered, and difficult to interpret (not shown). Thus, we attempted to independently predict 50 residue regions of Mrc1 in a 10-residue sliding window, in the presence or absence of Pol2 or either of its two halves (**Fig. S5A**). Using this multiparallel prediction strategy, we found one region that consistently had a high pLDDT score in the presence of Pol2^CTD^, but not Pol2^NTD^ (**Figs. S5B**,**C**). This region, MRC1^947–972^, was shown to have high sequence homology between fungal species (Naylor et al. 2009); we call this α-helix “N16” after that work. Intriguingly, we predict N16 to be nestled between a helix-loop-helix (HLH) motif on the Pol2^CTD^ surface that is situated adjacent to CMG, Mcm2 and Cdc45 (**Figs. 3A and S5D**). These subunits were shown by XL-MS to bind central regions of Mrc1 as well as more C terminal regions (Baretić et al. 2020) (**Fig. 2A**). In that report, Mrc1^917^ was crosslinked to Cdc45 and Mrc1^937^ was crosslinked to Mcm2, which would ideally position N16 for binding the nearest HLH of Pol2^CTD^, matching our prediction (**Fig. S5D**). Mrc1^752–end^ was previously shown by co-IP to bind only weakly to Pol2^CTD^ compared to more central regions of Mrc1 (Lou et al. 2008). However, aside from the aforementioned XL-MS studies that are odds with this, we also note that N16 was recently shown in CryoEM studies by the same group to bind to Pol ε^CTD^ at the meeting where we also first presented these data, validating our predicted model (Jones et al. 2025). We further predicted the existence of a salt bridge between Mrc1^K954^ and Pol2^D1955^ that stabilizes this interaction (**Figs. 3A, S5C, and S6**). These interacting residues are each highly conserved among yeast species, indicating the possibility that this Mrc1:Pol2^CTD^ interaction represents a conserved structural module (see MSA in **Fig. S6**).

**Figure 3.**
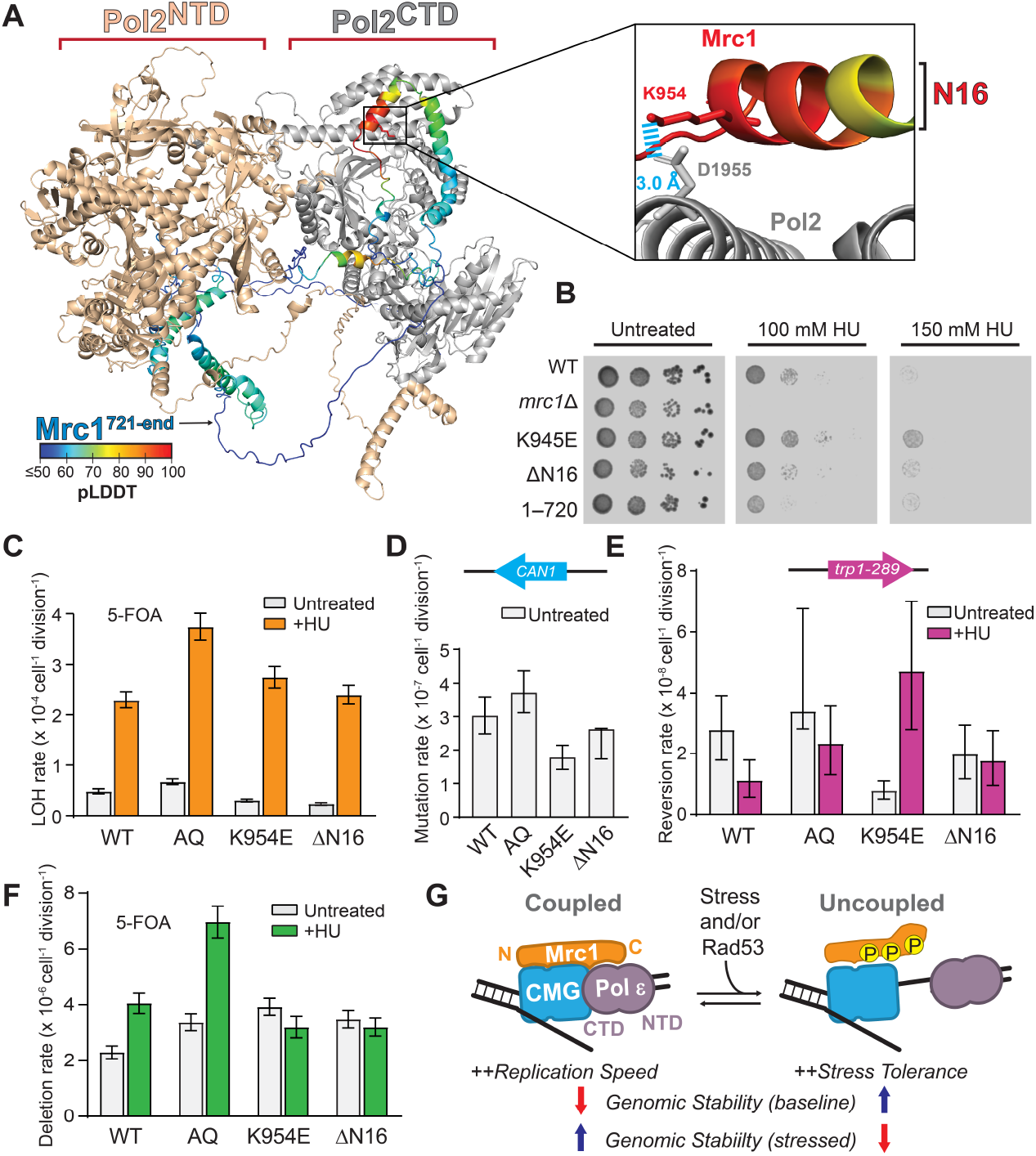
Forced helicase-polymerase uncoupling by disruption of a predicted Mrc1-Pol2 interaction confers higher baseline genomic stability and stress tolerance at the cost of stress-induced genomic instability. **A)** Predicted interaction of Mrc1^721–end^ with Pol2 highlighting a predicted salt bridge between Mrc1^K954^ and Pol2^D1955^. Pol2^CTD^ and Pol2^NTD^ are colored gray and beige, respectively. The pLDDT heat map used to color Mrc1^721–end^ is taken from the average of multiparallel AF2 runs (see Fig. S5). **B)** Survival assay of the indicated strains in the presence of the indicated HU concentrations. **C)** LOH rates at Chr. 4 in diploids (same assay as in Fig. 2C), **D)** Forward mutation rates at *CAN1* in haploids, **E)** Reversion rates of *trp1-289* in haploids, and **F)** Intrachromosomal deletion rates in haploids (same assay as in Fig. 2D) are shown for the indicated strains. Bars represent the mean mutation rate ± 95% CI; significance was inferred where CIs do not overlap, corresponding to *p* < 0.05. The full mutation rate dataset is provided in Fig. S9. **G)** Mrc1 controls fork coupling, genome stability, and stress tolerance. Uncoupling leads to flexibility and genomic stability in the baseline state but fragility in the stressed state, allowing enhanced stress tolerance at the cost of genomic instability. Conversely, a more rigid coupling of the fork that favors efficient progression through S-phase may lead to a slightly less stable genome overall in normal conditions, but in which genomic stability is prioritized under the existential threat of stress.

### Mrc1-Pol2 coupling: balancing baseline stability against stress-induced fragility

To test whether N16 and/or its salt bridge interaction with Pol2^CTD^ is important for the function of Mrc1, we made cells with *mrc1* bearing a charge-flip mutant (*mrc1*^K954E^) and cells with N16 deleted (*mrc1*^ΔN16^). In these experiments, we use *mrc1*^AQ^ as a comparative baseline for replication checkpoint deficiency that is less drastic than *mrc1*Δ. The results were particularly surprising for the charge-flip mutant in terms of growth on HU: we saw that K954E resulted in markedly increased HU tolerance (**Figs. 3B** and **S7**). Interestingly, using markers to measure LOH at two different chromosomal regions, we saw that both *mrc1*^K954E^ and *mrc1*^ΔN16^ exhibited significantly lower spontaneous LOH rates than WT cells (**Figs. 3C** and **S9B**) but showed similar or higher LOH rates than WT cells under HU stress (**Fig. 3C**). This trend—significantly decreased baseline mutation and increased perturbed mutation relative to wild-type for both the *mrc1*^K954E^ and *mrc1*^ΔN16^ strains—was also observed in haploid-based mutation assays, including those that measure point mutation rates and mutation reversion rates (**Figs. 3D, 3E, S9D**, and **S9E**). Conversely, we observed that *mrc1*^K954E^ and *mrc1*^ΔN16^ each displayed deletion rates that were significantly higher than WT cells at baseline, and that were significantly lower than WT cells under HU stress (**Fig. 3F**), mirroring the opposing trends that we saw with *mrc1*^1-946^ in **Fig. 2D**. Thus, disruption of the Mrc1:Pol2 interface appears to result in unique patterns of genetic instability that likely reflect differential replication-associated error events and repair pathways. We note that S965 is the only SQ/TQ residue on *mrc1*^947-end^, the region predicted to bind Pol2, and is adjacent to N16 (**Figs. 2A** and **S6A**), potentially making this residue key for checkpoint directed regulation of CMG:Pol ε coupling.

Taken together, our results are consistent with a model in which mrc1^1-720^ causes acute replication stress since it lacks much of the region required for robust checkpoint response (Naylor et al. 2009) and lacks the ability to slow runaway CMG down via Rad53 phosphorylation of the Mrc1 CTER (McClure and Diffley 2021). *mrc1*^1-946^ largely retains these abilities while remaining deficient in CMG-Pol ε coupling, resulting in increased HU tolerance but overall higher instability, likely due to impaired Mrc1 function including loss of many residues identified to be critical to direct CMG slowdown by Rad53 (McClure and Diffley 2021). Indeed, mutations in N16 (residues 947–972), including the K954E charge-flip mutation that specifically disrupts the Mrc1:Pol2 interaction, appear to confer a more controlled “forced uncoupling” phenotype, changing the balance of genome instability outcomes and suggesting a redistribution between replication-associated error events and repair outcomes following fork perturbation.

The pathways controlling these disparate mutation patterns remain to be characterized. While we do not directly assess fork remodeling, these data are consistent with a model in which disruption of the Mrc1:Pol2 interaction shifts the fork toward repair and/or remodeling pathways that favor replisome flexibility. In its coupled state, forks prioritize replication speed and stress resilience through enzymatic coordination. Disruption of the Mrc1:Pol2 tether forces the fork into a high-plasticity mode: while this state potentially excels at navigating spontaneous lesions through error-free remodeling, it lacks the structural reinforcement to prevent catastrophic collapse and LOH when the replication machinery is severely challenged by exogenous stress. We propose that Mrc1 directly mediates this delicate balance in a way that is tightly regulated by the constant feedback loop between the replisome and the checkpoint apparatus (**Fig. 3G**).

## MATERIALS AND METHODS

### Strains, plasmids, and oligonoucleotides

The base strain used for constructing overexpression strains is OY01. Haploid and diploid strains for mutation assays were based on JAY2087 (McLaughlin et al. 2020) and JAY3271. Construction of strains is described in Supplementary Materials. All strains, plasmids, and oligonucleotides generated for this study are listed in **Table S1 and footnotes and Table S2**.

### Protein purification

The purification of CMG, CMG-S6, Pol ε, Replication Factor C (RFC), PCNA, Sfp, and MTC (Mrc1, Tof1, Csm3) has been previously described (Schauer et al. 2017; Wasserman et al. 2019). The construction and purification of the Mrc1^AQ^, Tof1, Csm3 complex, CMG^555^, Pol ε^555^ is described in Supplementary Methods. Purity of protein complexes was assessed by coomassie stained SDS-PAGE, and all enzymes were assessed for activity *in vitro* before use in assays.

### Replication Uncoupling Assay

Replication uncoupling assays described in Results and Discussion used a modified version of primer extension assays on forked templates that have been described in detail (Schauer et al. 2017). Reaction volumes were 25 μL and contained 1.5 nM 2.8 kb GS501 forked template annealed to a 5’-^32^P-labeled primer. Reactions contained 25 mM Tris-OAc pH 7.5, 5% (v/v) glycerol, 50 μg/mL BSA, 2 mM TCEP, 3 mM DTT, 10 mM Mg-OAc, 50 mM K glutamate, 0.1 mM EDTA and were carried out at 30°C. DNA was first loaded with 30 nM CMG and 0.1 mM ATP for 10 min, followed by 20 nM Pol ε, 5 nM RFC, and 40 nM PCNA in the presence or absence of the indicated Mrc1-Tof1-Csm3 complex at 60 nM for 5 min. Reactions were initiated by the addition of 5 mM ATP, 120 μM dATP and dCTP, and 200 nM RPA, allowed to proceed for indicated uncoupling times (Δt), followed by addition of 120 μM dTTP and dGTP. Reactions were stopped with a solution containing 1% SDS and 40 mM EDTA. Reactions were run on a 1.2% alkaline agarose gel and visualized by phosphorimaging.

### Single-molecule FRET Replication Uncoupling Assay

smFRET assays used the same fork as the biochemical assays but it was 5’-biotinylated on the lagging strand and used a shorter template (209 bp). Reactions followed the biochemical uncoupling assay apart from withholding dTTP and dGTP at the last step to prevent replisome runoff from template and additionally contained an oxygen scavenging mixture (25 nM PCD, 2.5 mM PCA, 1 mM Trolox, 1 mM cyclooctatetraene, and 1 mM 4-nitrobenzyoyl alcohol). Reactions were imaged on a homebuilt prism-based TIRF microscope based on a Nikon Eclipse Ti2 with a SR Plan Apo IR 60X objective. 532 nm and 640 nm wavelength lasers were used for alternating laser excitation to selectively excite LD555 and LD655, respectively. Data was analyzed with SPARTAN using semi-automatic filtering of traces.

### Western Blot

Primary antibodies: Rabbit anti-Rad53 (Abcam AB104232) at 1:2000, or mouse anti-α-tubulin (ABM # G094) at 1:2000 for loading control. Secondary antibodies: DyLight goat anti-mouse 680 nm, or goat anti-rabbit 680 nm, at 1:15,000. See Supplementary Methods for full details.

### Serial spot dilution assays

Yeast spot assays were performed with yeast harvested in exponential phase. Pellets were resuspended in sterile water and serially diluted to 10^5^, 10^4^, and 10^3^ cells/mL with sterile water; spots were 5 μL.

### Genetic mutation assays

Strains were grown and treated as indicated in Supplementary Methods. Yeast were plated on indicated medium and incubated for 3 days at 30°C. Colonies were counted manually, and counts were used to calculate maximum likelihood mutation rates and 95% confidence intervals.

### Multiparallel predictive structural modeling

AF2 predictions of Mrc1 or Mrc1^721–end^ were performed in 50 residue windows sliding by 10 residues across the length of the protein, in the presence or absence of Pol2, Pol2^NTD^, or Pol2^CTD^ (see Fig. S6) using Colabfold running on a local cluster (Mirdita et al. 2022). Sliding averages of the pLDDT scores and interatomic distances of the top models (out of 5 models per run) were calculated with python scripts.

## ACKNOWLEDGMENTS

The funding for this work was provided by the NIH (R35GM147105) to G.D.S. and (R35GM119788) to J.L.A. V.M.L. was supported by an NIH fellowship (T34GM140958). High performance computer time and resources were provided by Colorado State University through the Data Science Research Institute. We thank Mattias Mihelich and Jackson Whitted for assistance with genetic experiments, and Alisa Shaw for useful guidance.

## Author Contributions

G.D.S. administered the project, G.D.S. and J.L.A. conceived the study, and G.D.S., J.A.S., and J.L.A. designed the methodology. All authors collected and/or analyzed the data. A.J.T, J.L.A., and G.D.S. wrote the manuscript.

