## Supplementary Info for "Mechanical tuning of replication stress tolerance and genomic stability through the checkpoint mediator Mrc1"

**SUPPLEMENTARY INFORMATION.** Timmerman et al., 2026

### Supplementary Figures

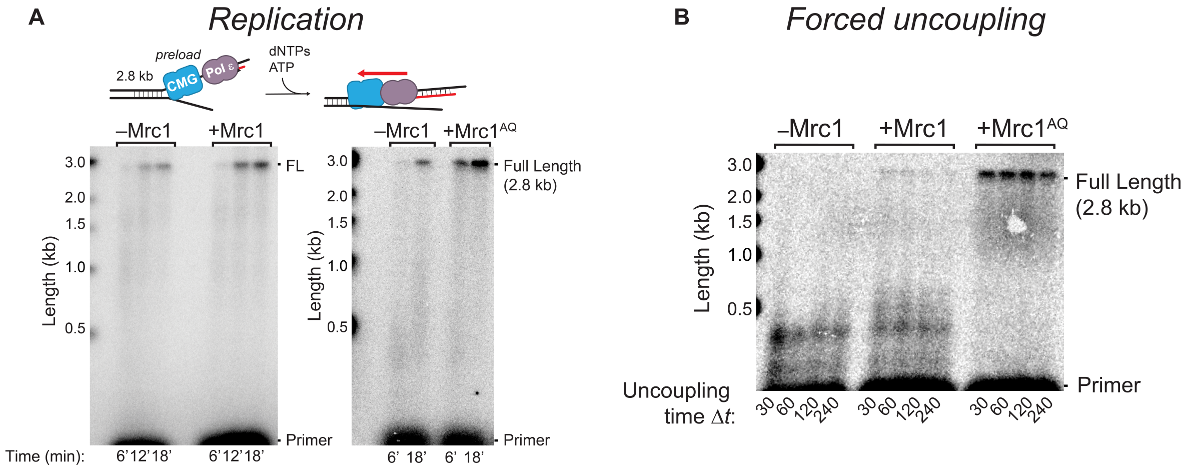


**Figure S1. Mrc1$^{AQ}$ enhances replication speed similar to WT Mrc1 A)** Mrc1 and Mrc1$^{AQ}$ stimulate replication speed by ~1.5 fold. Regular forked 2.8 kb replication reactions were performed as previously described (Schauer et al. 2017) in the presence of MTC or M$^{AQ}$TC and stopped at the indicated times; fork speed determination was made by comparing integrated full length product intensities at respective timepoints via densitometry analysis. Reactions also contained RFC, PCNA, and RPA (not shown for clarity). **B)** Additional example of uncoupling experiment used in the quantification for Fig. 1C, increasing contrast relative to Fig. 1B so the full-length product in the presence of WT Mrc1 can be seen more readily.

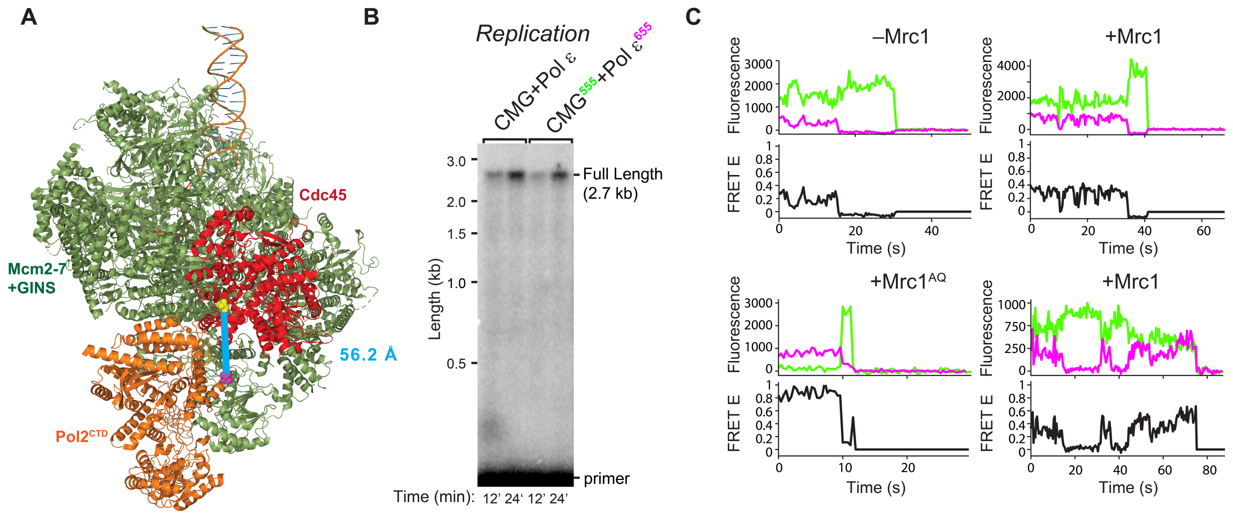


**Figure S2. smFRET construct design, labeled protein activity, and example smFRET traces. Validation. A)** Molecular model of CMG-Pol ε labeling scheme, with proteins colored as indicated. Using additional 11 residue S6 peptides (which are not modeled here), CMG was labeled with LD555 at the CTER of Cdc45 (yellow spheres), and Pol2 was labeled at its CTER (magenta spheres). A FRET=1 value indicating < ~3 nm inter-dye distance is possible: the distance between the residues shown is ~5.6 nm (blue line), there are two additional S6 peptides at each C-terminus, and the distance of the dyes and their linkers is ~ 2nm each. Model adapted from PDBID:8KG6 (Xu et al. 2023) by removing Ctf4, Tof1, and Csm3. **B)** Dye labeling does not affect activity of either CMG or Pol ε. Regular replication reactions of CMG and Pol ε (left) vs CMG$^{555}$ and Pol ε$^{655}$ (right) are shown. **C)** Representative smFRET traces are shown for the indicated conditions. Donor (555): green; Acceptor (655): magenta; FRET E: black. For each condition, an acceptor-then-donor photobleaching event is shown to demonstrate canonical FRET. An additional trace for the +Mrc1 condition is shown (bottom right) to demonstrate the dynamic behavior observed in this condition.

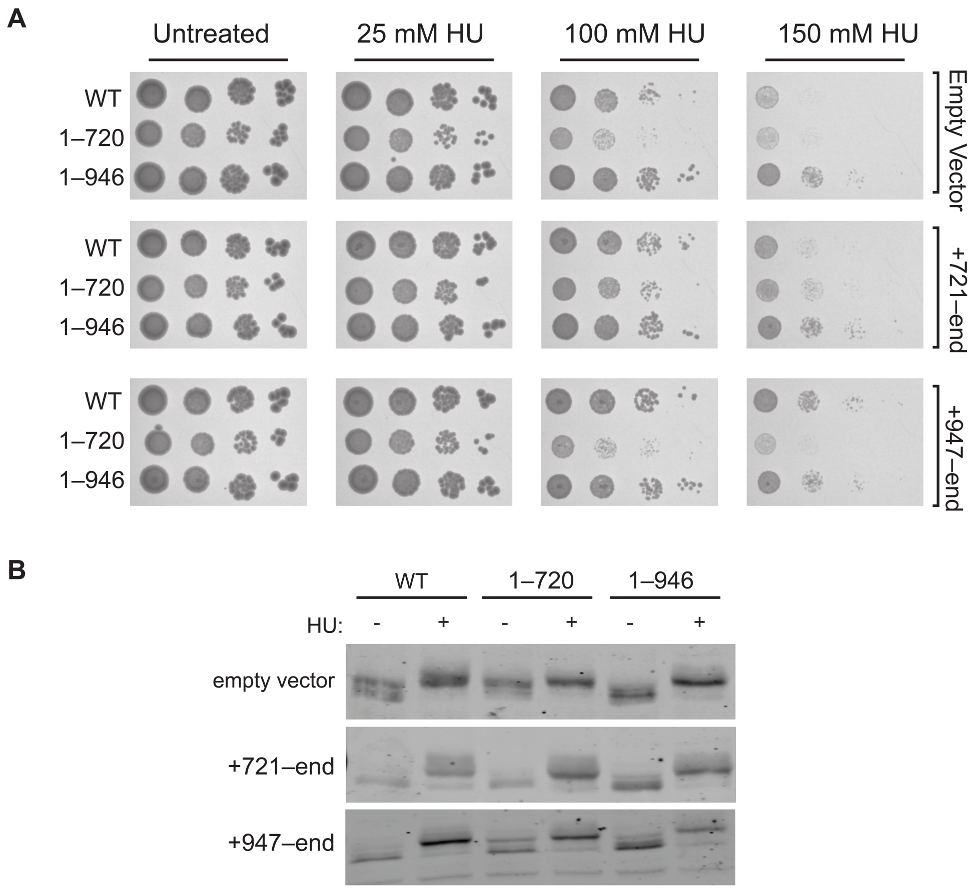


**Figure S3. Increased survival from Mrc1 fragment rescue is not likely due to increased Rad53 activity. A)** Biological repeat of survival spot assays shown in Fig. 2B. Cells bearing the indicated *mrc1* truncations are shown in response to 150 mM HU stress where indicated. Where indicated, cells were transformed with single-copy CEN plasmids expressing the specified *mrc1* C-terminal fragment. **B)** Rad53 Western blot for the indicated conditions.

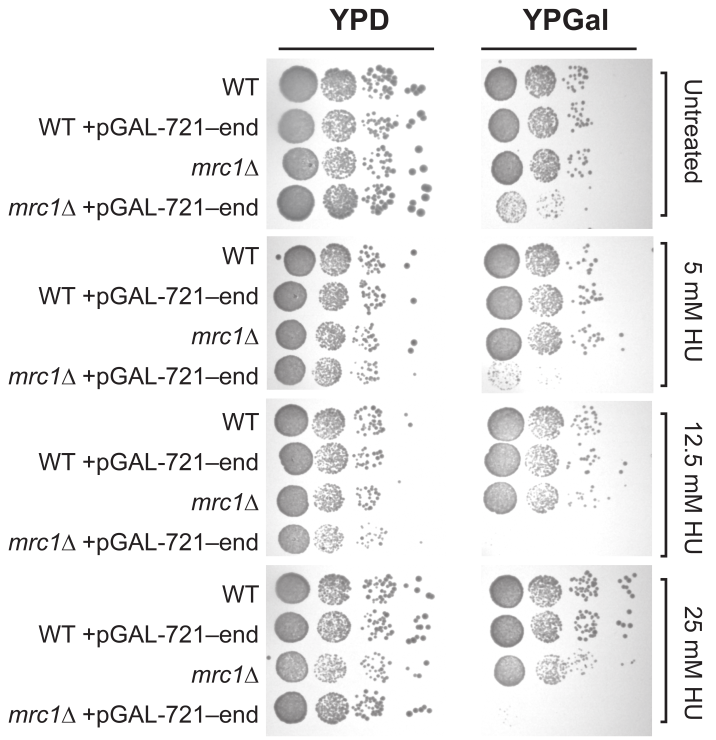


**Figure S4. Overexpression of *mrc1*$^{721\text{-end}}$ causes lethality and extreme HU sensitivity in strains lacking MRC1.** Survival spot assays are shown for the indicated strains at the indicated HU levels. Where noted, strains were integrated with pGAL expression cassettes for Gal-inducible overexpression of the indicated construct.

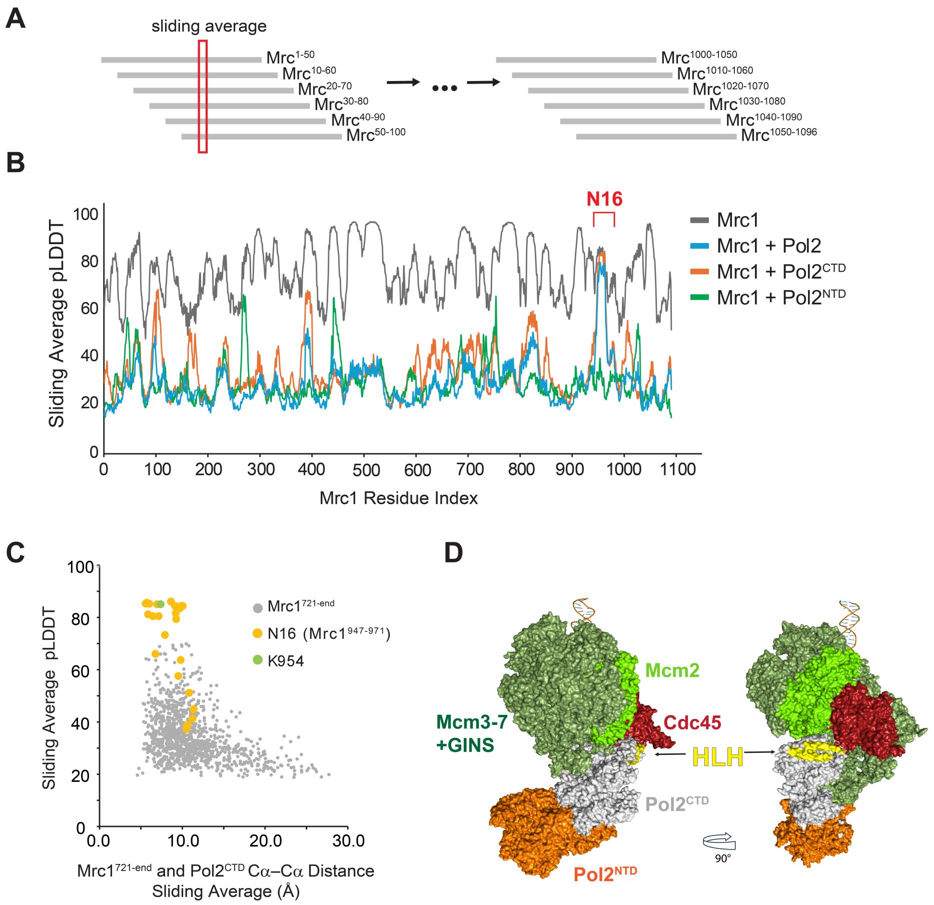


**Figure S5. Multiparallel predictive modeling identifies Mrc1$^{N16}$ as an interaction partner with Pol2$^{CTD}$. A)** Schematic of prediction algorithm. **B)** Sliding average AlphaFold 2 (AF2) pLDDT of the indicated protein complexes is shown. AF2 predictions were performed in 50 residue windows, sliding by 10 residues. The plot represents an average pLDDT of the top models from all windows covering each residue and the pLDDT heatmap of Mrc1$^{CTD}$ in Figure 3 is colored according to this sliding average. The N16 alpha helix (residues 947-971) is indicated, which was predicted to bind the CTD, but not NTD, of Pol2. **C)** Sliding average pLDDT vs. Sliding average Cα-Cα distance between Mrc1$^{721\text{-}end}$ and Pol2$^{CTD}$. For each Mrc1 residue along the sliding window of AF2 runs, the interatomic Cα-Cα distance to the nearest Pol2 Cα was calculated for the top scoring model coordinates and averaged across all windows for each residue. N16 and K954 are highlighted, demonstrating the region predicted to have the shortest Mrc1-Pol2 distance concomitant with high prediction confidence. **D)** Model of the Helix-Loop-Helix (HLH; yellow) of Pol2$^{CTD}$ (residues 1948-2021) in context of CMGE. The cryoEM structure of Pol ε (PDB:6WJB) (Yuan et al. 2020) was aligned to the cryoEM structure of CMGE, which partially contains Pol2$^{CTD}$ (PDB:8KG6) (Xu et al. 2023). Tof1, Csm3, and Ctf4 were removed for clarity.

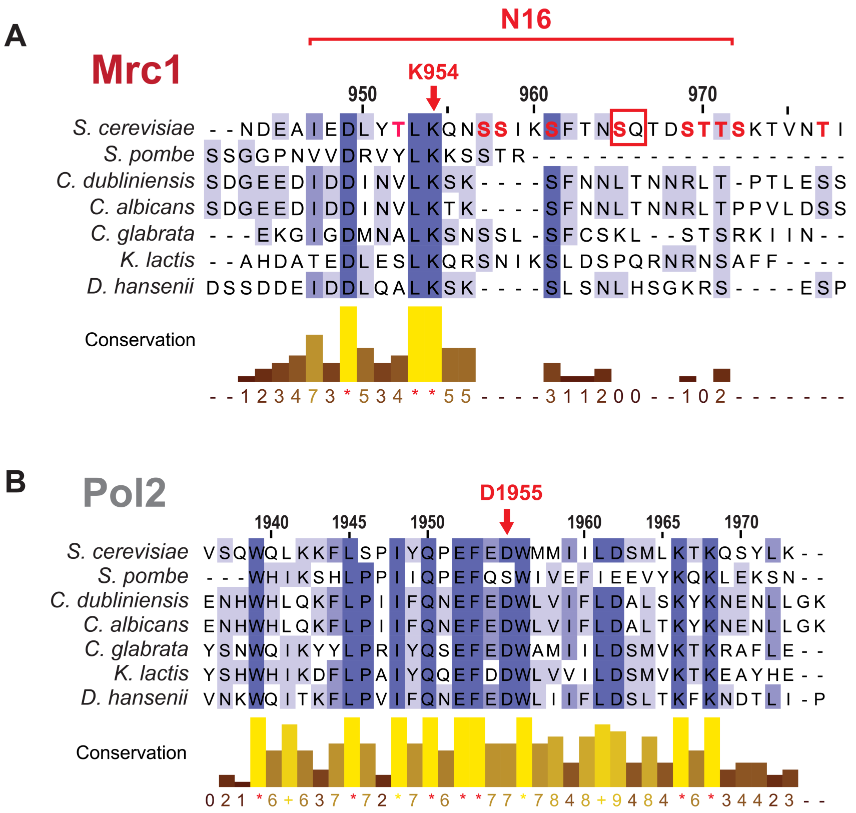


**Figure S6. Multiple sequence alignment of predicted Pol2 and Mrc1 interacting domains.** Amino acid sequences of **A)** *Sc*MRC1 and **B)** *Sc*POL2, as well as their indicated fungal homologs, were obtained from the UniProt database. These sequences were aligned via Jalview (Waterhouse et al. 2009) using the MAFFT algorithm (Katoh and Standley 2013). N16 and the predicted Mrc1-Pol2 interaction salt bridge residues are indicated. Residues which can be phosphorylated on Mrc1 are colored red, and the SQ/TQ motif residue within N16 is highlighted with a red box.

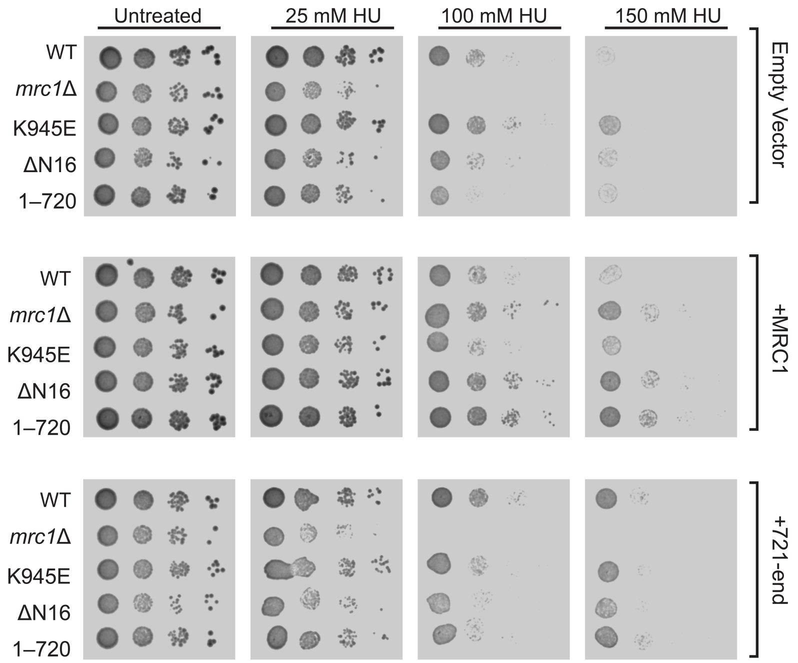


**Figure S7. Further characterization of strains and near-native expression rescue by MRC1 and *mrc1*$^{721\text{-}end}$.** Survival spot assays are shown for the indicated strains at the indicated HU levels. Where noted, strains were transformed with CEN expression vectors for the indicated construct.

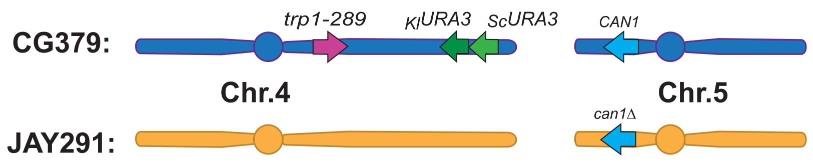


**Figure S8. Schematic representation of mutation reporter markers**
The upper blue chromosomes represent the mutation detection reporters present in the *MAT*$\alpha$ CG379-isogenic haploid strains carrying the various mutations in *MRC1*. When grown as haploids, these strains were used to measure the rates of interstitial deletion of the tandem *URA3* markers on the right arm of Chr. 4, reversion to Trp+ of the *trp1-289* mutation, and forward mutation in the *CAN1* gene on the left arm of Chr. 5. Note that the tandem *URA3* cassette on Chr. 4 is flanked by ~500 bp directly-oriented repeats that serve as substrates for homology-mediated deletion (schematic in **Fig. 2D**). These repeats are ~5 kb away from each other. The lower orange chromosomes represent the configuration of Chr. 4 and Chr. 5 in the *MATa* JAY291-isogenic haploid strain AT17 (*mrc1Δ*). When crossed to the CG379-derived haploid strains carrying the various mutations in *MRC1*, the resulting hybrid diploids were used to measure the rates of LOH on the right arm of Chr. 4 and the left arm of Chr. 5. Circles represent the approximate positions of centromeres. Additional details provided in **Table S1** and footnotes.

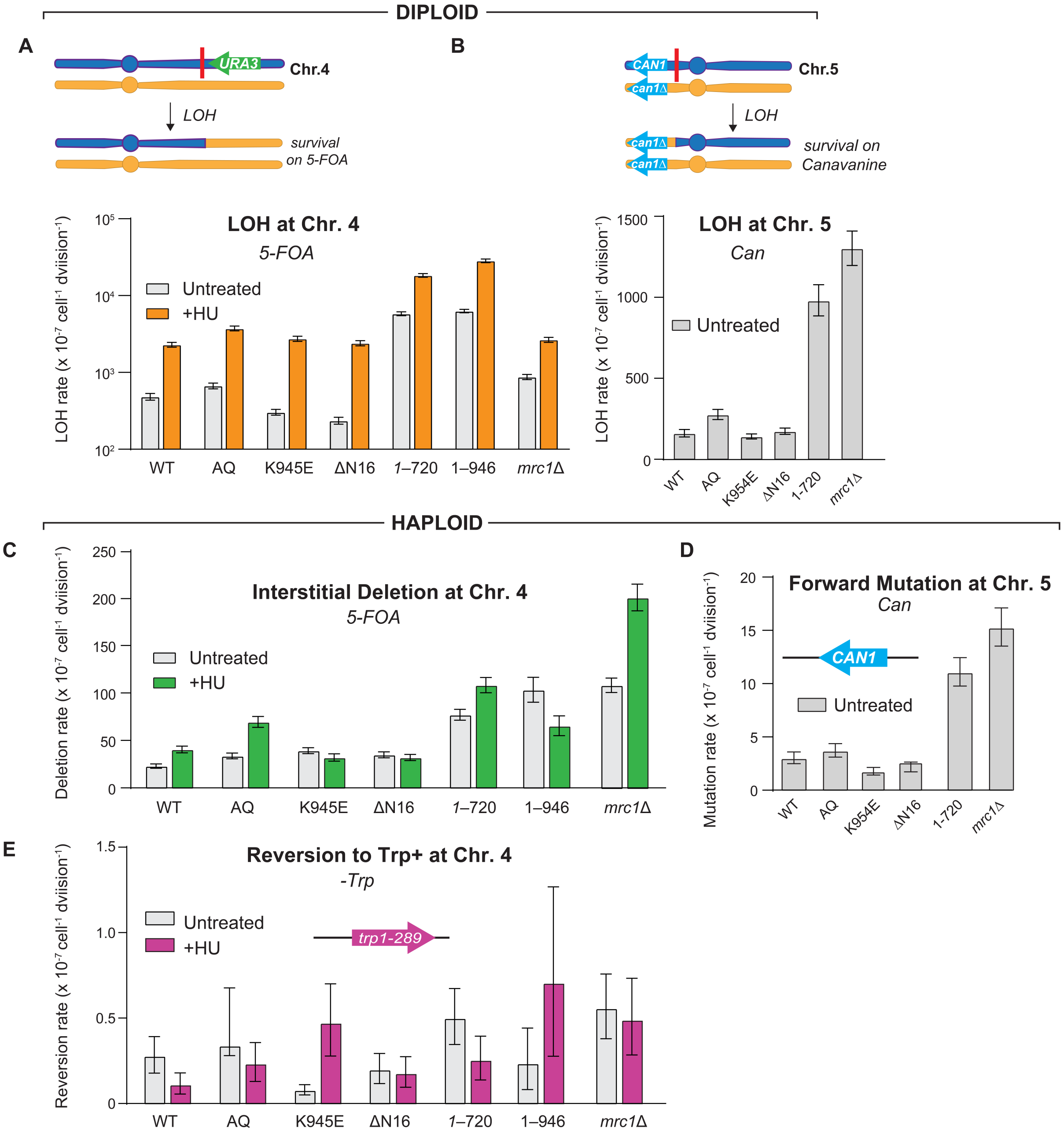


**Figure S9. Genetic assays summary**
Plots showing the entire dataset of all mutation rates and 95% C.I. determined through this study. Upper plots **A)** and **B)** show the results of the indicated LOH assays in diploid strains. Lower plots **C)**, **D)**, and **E)** show the results of indicated mutation classes measured in haploids. Schematics above each plot show the relevant arrangement of mutation detection markers in the assay strains as in **Fig. S8**, as well as the respective selective media used. Subsets of these data points are duplicated in **Fig. 2** and **Fig. 3** of the main manuscript. Additional details are provided in **Table S1** and footnotes.

### Supplementary Methods and Text

*Yeast strain construction*

Strains used in this study, listed in **Table S1**, were generated from one of three base strains: OY01, JAY2087, or JAY3271. In the cases of AES10, AT18, AT19, AT20, AT21, and AT23, CRISPR-mediated mutagenesis was used as described (Akhmetov et al. 2018 for AES10, AES11, and AES12; Anand et al. 2017 for AT18, AT19, AT20, AT21, and AT23). For all others, traditional yeast genetic manipulation was performed using standard replacement of genes with various marker cassettes. All yeast cultures were grown in YPG/Gal media at 30°C. To knock out genes or truncate them, the coding sequence was replaced via a donor containing a selectable marker flanked by homology for the target gene UTR or coding sequence. To integrate a gene whose expression was driven by GAL1/10, the gene was introduced to a pRS40x-GAL series vector via restriction cloning. The vector was then linearized within the nutritional marker via restriction digest and transformed into LiAc-competent yeast via heat shock protocol. To generate hybrid diploid yeast for LOH assays, *MATa* and *MAT$\alpha$lpha* haploid parent strains were plated on YPD after mixing in sterile ddH2O. Resultant patches were streaked to single colonies, which were screened for diploids by plating on selection according to the expected phenotype of the hybrid diploids. Selected diploids were used as described in LOH assays. Additional strain information including genetic backgrounds and their respective uses for the various experiments described in this study are provided as footnotes to **Table S1**.

*Protein Purification and Labeling*

Yeast protein complexes (CMG, CMG-S6, Pol $\varepsilon$, MTC, RFC, PCNA, RPA) were purified from yeast overexpression strains by chromatography as previously described (Georgescu et al. 2014, 2015; Langston et al. 2017; Wasserman et al. 2019; Lewis et al. 2020). Fluorophore labeling reactions were performed enzymatically with Sfp targeting a genetically encoded S6 fusion tag (Zhou et al. 2007) using 1:2:4 ratio of protein-S6:Sfp enzyme:CoA-fluorophore for 16 hours at 4°C. Free dye was removed from the reaction via ZebaSpin 2 mL desalting columns (Thermo #89889). Labeling efficiency was assessed by fluorescence scan on a multiwavelength scanner (GE Typhoon FLA 9500) prior to Coomassie staining of SDS-PAGE gels.

*Mrc1$^{AQ}$-Tof1-Csm3 purification*

Purification of Mrc1$^{AQ}$-Tof1-Csm3 closely followed previously published purification of MfTC (Lewis et al. 2020). Briefly, cultures of Mrc1$^{AQ}$-3xFLAG-Tof1-Csm3 overexpression strain (described in **Table S1**) were grown in YP-glucose media to log-phase and then switched into YP-glycerol media. Cultures were again grown to log-phase and then induced by addition of 2% galactose for 6 hours. Cells were pelleted at 2,000 x g for 10 minutes, and cell pellets were resuspended with 25 mL of lysis buffer: wash buffer (10% glycerol, 300 mM Potassium Glutamate, 50 mM Hepes pH 7.5, 1 mM EDTA pH 8.0, 0.5 mM DTT) supplemented with 100 µL protease inhibitor cocktail (Sigma #P8215) per 20 g dry pellet). Resuspended pellets were then dripped into liquid nitrogen and stored at -80°C. Frozen pellet droplets were mechanically lysed in a Spex CG-500 Freezer/Mill (12 cycles, 15 cps, 2-minute run), which was subsequently thawed in the cold room with gentle stirring and 25 mL additional wash buffer. Lysates were clarified by centrifugation at 19,000 x g for 1 hour, then again for 30 minutes, before loading on equilibrated anti-FLAG M2 affinity resin (Sigma #A2220). The resin was washed with 40 column volumes of wash buffer, and then protein was eluted using 6 CV of wash buffer containing 0.5 mg/mL 3xFLAG peptide (GLPBio #GP10149). Purity was assessed by Coomassie-stained SDS-PAGE, and yield was quantified by BioRad protein quantification assay. Mrc1$^{AQ}$ was assayed by primer extension assay to be similarly active to WT Mrc1 in terms of fork speed enhancement (**Fig. S2B**).

*CMG$^{555}$ purification*
CMG-S6 (CMG with Cdc45-S6-3xFLAG) was purified as previously described (Wasserman et al. 2019) and subsequently labeled with CoA-LD555 (Lumidyne Technologies) as described above. Labeling efficiency was determined to be > 90%; replication-coupled unwinding activity was assayed by replication assay and determined to be indistinguishable from WT in terms of activity (**Fig. S2B**).

*Pol $\varepsilon^{655}$ purification*
Pol ε was purified as previously described (Georgescu et al. 2015) with the exception of adding an S6 tag between the C-terminal domain of Pol2 and its 3XFLAG tag. Pol $\varepsilon^{655}$ was purified as described for WT Pol ε, employing tandem flag and heparin column purification (Georgescu et al. 2014). The prep was labeled with CoA-LD655 (Lumidyne Technologies); labeling efficiency was > 90% and intrinsic activity in reconstituted replication reactions was unaltered (**Fig. S2B**).

*Single-molecule FRET*
Single-molecule FRET experiments were performed on a homebuilt prism-based total internal reflection fluorescence (TIRF) setup based on a Nikon Ti2-E inverted fluorescence microscope. Flow cells were constructed from Quartz slides (G. Finkenbeiner, Inc.) and an ~80 uM adhesive gasket (ARcare 90445). Slides were PEGylated with mPEG and bioPEG (Laysan Bio) and incubated with Neutravidin (Sigma) as described (Joo and Ha 2012). Biotin-conjugated DNA was surface-tethered at 50 pM. General assay conditions were identical to those used in the biochemical uncoupling assays and additionally included the protocatechuic acid/protocatechuate-3,4-dioxygenase (PCA/PCD) oxygen scavenging system (Aitken et al. 2008) with 40 nM PCD, 2.5 mM PCA, 1 mM Trolox, 1 mM cyclooctatetraene, and 1 mM 4-nitrobenzyl alcohol to prevent photobleaching and/or blinking. Continuous wave lasers (488 nm/100 mW, 532 nm/80 mW, and 640 nm/75 mW; Coherent OBIS) were used continuously or alternately by exciting 555 and 655 dyes every other frame, a.k.a. Alternating Laser Excitation (ALEX). Emission was collected with a 60X Plan Apo TIRF water immersion objective, sent through an image splitter (Cairn) with a 560 nM longpass (LP) dichroic mirror (Chroma T560lpxr-UF1) and a 640 nM LP dichroic (Chroma T640lpxr-UF2). The 488 and 555 dye emissions were filtered through cleanup filters (Chroma ET525/50m and 585/65m, respectively). Spectrally filtered images were projected onto the focal plane of three scientific complementary metal-oxide Semiconductor (sCMOS) cameras (ORCA Fusion, Hamamatsu), and movies were recorded at a rate of 200 ms per frame. The data was parsed using custom Python scripts and analyzed using SPARTAN (Juette et al. 2016). All FRET trajectories were filtered to remove post-photobleaching data, and ALEX was used to select traces with FRET pairs to eliminate choosing donor-only peaks and other potential artifacts.

*Tandem Mass Spectrometry (MS/MS)*
To understand whether yeast-purified Mrc1 carries basal phosphorylation acquired *in vivo* during normal growth and purification, we analyzed a sample of purified Mrc1 by tandem mass spectrometry (MS/MS) at the Proteomics Core Facility at University of Colorado, Anschutz. Cultures of the yeast strains for purification of WT Mrc1 were grown as for our regular purification, starting in YP-glucose media to log-phase and then switch to YP-glycerol media and again grown to log-phase. Cultures were then induced with 2% galactose for 6 hours. Cells were pelleted and lysed as described above, with the addition of 1 µM microcystin-LR (Enzo #ALX-350-012) and 1 phosSTOP tablet per 50 mL buffer (Sigma #4906845001). Purification of MTC was carried out as described above, with the same concentration of phosphatase inhibitors added to lysis and initial chromatography buffers. MTC was run on SDS-PAGE and stained with Coomassie. Mrc1 bands were cut out of the gel and flash-frozen before subsequent analysis. We estimated the abundance of phosphorylated residues semi-quantitatively by

confirming mean intensities of trypsinized peptides between two separate replicates. We identified phosphorylation at 13 residues (S112, S121, S144, S367, T373, S388, S434, S605, S607, S609, S622, S924, and S1010), including a single SQ/TQ motif (S434), indicating that SQ/TQ residues are phosphorylated in WT cells lacking chemical stress, though undergoing potential stress associated with purification. Despite an overall sequence coverage of 70.3%, coverage across regions containing potential phosphosites was uneven, and S434 was the only SQ/TQ site detected in the dataset, regardless of phosphorylation state assignment. This may reflect lower detectability of peptides from these regions, which can be influenced by properties such as intrinsic disorder and reduced ionization or fragmentation efficiency in mass spectrometry. Together, these data indicate that phosphorylated species of WT Mrc1 are present in the purified sample under the conditions tested.

*Western blots*
Yeast strains were grown to log phase in appropriate medium at 30°C. Then, α-factor (EZBiolab Custom Peptide (PT0803180601)) was added to a final concentration of 0.1 µg/mL to arrest cells at the G1/S interphase for 3 hours. After the 3 hours arrest, cells were washed three times with sterile water before being resuspended in fresh, pre-warmed medium. Resuspended cells were then treated with addition of 150 mM hydroxyurea for 1 hour at 30°C. After 1 hour, 1.5 mL of these cultures were pelleted and flash frozen in liquid nitrogen for storage at -80°C until lysis. The lysis was performed via an adapted method as described (Cox et al. 1997). Briefly, frozen pellets were thawed on ice with 300 µL of TCA lysis buffer (10 mM Tris-HCl, pH 8.0; 10% trichloroacetic acid (TCA); 25 mM $NH_4OAc$; 1 mM EDTA) and 100 µL 0.5 mm glass beads. Samples were vortexed for 1 minute, then kept on ice for 3 minutes, repeated five times. Lysates were then removed and pipetted into a new tube, with an additional 100 µL TCA lysis buffer wash of the glass beads, and then pelleted at 12,000 x g for 15 minutes at 4°C. Supernatant was removed, and pellets were resuspended in 40 uL resuspension solution (100 mM Tris-HCl, pH 11.0; 3% SDS) before being boiled for 5 minutes. After cooling at room temperature for a few minutes, samples were centrifuged again at 16,000 x g for 30 seconds, and the supernatant was moved to a fresh tube. Protein concentration was quantified using BioRad DC assay, after which a normalized amount of sample was added to gel loading buffer (250 mM Tris pH 6.8, 10% glycerol, 1% SDS, 10% β-mercaptoethanol, 0.005% bromophenol blue). Samples were run on 7.5% SDS-PAGE gels and subsequently transferred to nitrocellulose membranes. Membranes were blocked in 1% BSA in TBS (10 mM Tris pH 8.0, 150 mM NaCl). Antibody incubations and washes were in TBST (10 mM Tris pH 8.0, 150 mM NaCl, 0.1% Tween-20), with 1% BSA present in antibody incubations. Primary antibodies: rabbit anti-Rad53 (Abcam AB104232) at 1:2000, or mouse anti-α-tubulin (ABM # G094) at 1:2000 for a loading control. Secondary antibodies: DyLight goat anti-mouse 680 nm, or goat anti-rabbit 680 nm, at 1:15,000. Blots were detected on an Odyssey CLx infrared scanner.

*Genetic mutation assays and strain constructions*
Yeast strains were generated from the base haploid strains *MATαlpha* JAY2087 (McLaughlin et al. 2020), which contains multiple selectable markers used to quantify the rates of various types of mutation, and *MATa* JAY3271 which was used as the complimentary mating parent to generate hybrid diploids for LOH assays. JAY2087 haploids are isogenic with the CG379 genetic background (Morrison et al. 1991), and JAY3271 is isogenic with JAY291 (Argueso et al. 2009). See **Table S1** and footnotes and **Fig. S8**.

Mutation assays in haploid strains derived from JAY2087: Homology-driven interstitial / intrachromosomal deletion was measured via counter selection for resistance to 5-fluoroorotic acid (5-FOA) gained from the deletion of tandem *URA3* genes flanked by direct repeat sequences (McLaughlin et al. 2020). Forward mutation of *CAN1* was measured via counter selection for resistance to canavanine through inactivation of *CAN1*. Reversion of point mutations is measured via tryptophan prototrophy acquired by reversion of a nonsense point mutation in the *trp1-289* allele.

Mutation (LOH) assays in hybrid diploid strains: Haploids derived from JAY2087 were mated to AT17. LOH at the tandem *URA3* cassette inserted on the right arm of chromosome IV distal to *PLM2* was measured through counter selection for 5-FOA resistance. LOH at the *CAN1* locus on the left arm of chromosome V was measured through counter selection for canavanine resistance.

Strains were streaked on YP-glucose plates and grown at 30°C until isolated colonies formed, about three days for haploids and four days for diploids. For assays using haploid strains, the cultures were expanded in a second phase of growth by inoculating single colonies into liquid YP-glucose medium and grown for 22 hours, without or with 25 mM hydroxyurea where applicable. 1 mL of these cultures were pelleted; the pellets were washed once with sterile water, then spun down and resuspended once again in fresh sterile water. For LOH assays using diploids, strains were streaked on YP-glucose plates without or with 25 mM hydroxyurea where applicable. Colonies were then taken directly from the streaking plates and resuspended in sterile water. The resuspended yeast pellets were serially diluted by a factor of 10 up to $10^{-5}$ and plated as follows: For haploid assays, 150 µL of $10^{-5}$ on YPD, 150 µL of $10^{-1}$ on 5-FOA, 200 µL undiluted on canavanine, and 600 µL undiluted on tryptophan dropout; For diploid LOH assays, 100 µL of $10^{-4}$ on YPD, 50 µL of $10^{-1}$ on 5-FOA, and 25 µL undiluted on canavanine.

Multiple independent cultures were generated and plated for each *MRC1* genotype and HU exposure condition. After three days, colonies were counted from selective and permissive media, and counts were used to calculate mutation rates and 95% confidence intervals using the MSS Maximum Likelihood method with the FALCOR web application (https://lianglab.brocku.ca/FALCOR/ ; Hall et al. 2009).

*Analysis of spectrum of forward mutation in CAN1*
We carried out a preliminary PCR-based assessment of the qualitative classes of mutations associated with forward mutation to canavanine resistance in haploid strains. Specifically, we differentiated between short mutations (*i.e.*, point mutations and small frameshift insertion/deletions) versus larger deletions (~50+ bp) within the *CAN1* gene, using an approach adapted from described previously (Xie et al. 2001).

Haploid strains AT13 (WT; *MRC1::HphMX6*), AT14 (*mrc1(AQ)::HphMX6*), AT15 (*mrc1(1-720)::NatMX6*) and AT16 (*mrc1Δ::NatMX6*) were streaked to single colonies in YP-glucose plates. After 3 days incubation at 30C, ~30 independent single colonies from each strain were picked with a toothpick and patched (3 cm x 3 cm) to agar media containing canavanine. After 3 days incubation at 30°C, one canavanine-resistant (CanR) colony/papillae was picked per patch to ensure mutational independence. Each independent CanR papillae was streaked in media containing canavanine to be purified to a single clone. Genomic DNA from each CanR clone was extracted and used in PCR with primers JAO270 and JAO273 to amplify across the *CAN1* gene. The 1,954 bp PCR product was then digested with *Taq*I-v2 (New England Biolabs; R0149S; T▼CGA) frequent cutter restriction endonuclease to generate shorter DNA fragments (8, 42, 162, 332, 671, and 749 bp), and then run on a 2% agarose gel for separation and assessment of the structure of *CAN1* in each CanR clone. DNA from the *CAN1* WT (CanS) parent strain was used as a control reference, in which the four longer fragments were clearly detectable (8 and 42 bp were not visible). In all cases, (WT [21 of 21 CanR clones], *mrc1(AQ)* [29 of 29], *mrc1(1-720)* [31 of 31], and *mrc1Δ* [31 of 31]), the *Taq*I-v2 restriction banding pattern of the *can1* mutations matched the pattern in the *CAN1* control. This result showed that spectrum of forward mutation in the *CAN1* gene in the three *MRC1* alleles analyzed was indistinguishable from wild type, consisting primarily of point mutations or small insertion/deletions that do significantly alter the size of DNA fragments at the resolution of this assay. This indicated that the quantitative ~5-fold increase in the rate CanR forward mutations measured in *mrc1Δ* (**Fig. S9**) was not due to a qualitative spectrum change, such as structural rearrangements. Instead, our results showed that the quantitative increase in CanR mutations in *mrc1Δ* is likely primarily due to a lower fidelity of DNA copying, leading to a higher rate of point nucleotide substitutions or small insertion/deletions qualitatively similar to that seen in wild type. For context, the mutational spectrum observed in this assay for some DNA replication mutants such as *rad27Δ* is characterized by a pronounced increase in the frequency of larger deletions (50+ bp) within the *CAN1* locus (Xie et al. 2001).

**TABLE S1: Strains used in this study**

| Strain Name | Genetic background | Genotype |
|---|---|---|
| OY01 | [a] W303 | *ade2-1 ura3-1 his3-11,15 trp1-1 leu2-3,112 can1-100 bar1Δ MATa pep4::KANMX6* |
| OY6 | [a] W303 | *ADE2::GAL-MRC1-3xFLAG ura3-1 HIS3::GAL-TOF1 trp1-1 LEU2::GAL-CSM3 can1-100 bar1Δ MATa pep4::KANMX6* |
| AES10 | [a] W303 | *ADE2::GAL-mrc1(AQ)-3xFLAG ura3-1 HIS3::GAL-TOF1 trp1-1 LEU2::GAL-CSM3 can1-100 bar1Δ MATa pep4::KANMX6* |
| AES13 | [a] W303 | *HIS3::GAL-PSF1, PSF2 ADE2::GAL-PSF3, SLD5 LEU2::GAL-MCM6, MCM7 TRP1::GAL-MCM3, MCM4 MET15::GAL-CDC45-S6-FLAG URA3::GAL-MCM2, HK/P-MCM5* |
| AT03 | [a] W303 | *ade2-1 ura3-1 his3-11,15 trp1-1 leu2-3,112 can1-100 bar1Δ MATa pep4::KANMX6 mrc1Δ::TRP1* |
| AT20 | [a] W303 | *ade2-1 ura3-1 his3-11,15 trp1-1 leu2-3,112 can1-100 bar1Δ MATa pep4::KANMX6 mrc1-K954E* |
| AT21 | [a] W303 | *ade2-1 ura3-1 his3-11,15 trp1-1 leu2-3,112 can1-100 bar1Δ MATa pep4::KANMX6 mrc1Δ(950-961)* |
| AT22 | [a] W303 | *ade2-1 ura3-1 his3-11,15 trp1-1 leu2-3,112 can1-100 bar1Δ MATa pep4::KANMX6 mrc1(1-720)::natMX6* |
| AT24 | [a] W303 | *ade2-1 ura3-1 his3-11,15 trp1-1 leu2-3,112 can1-100 bar1Δ MATa pep4::KANMX6 mrc1(1-946)::natMX6* |
| AT25 | [a] W303 | *ade2-1 URA3::GAL-mrc1Δ(1-720)-3xFLAG-NLS his3-11,15 trp1-1 leu2-3,112 can1-100 bar1Δ MATa pep4::KANMX6* |
| AT26 | [a] W303 | *ade2-1 URA3::GAL-mrc1Δ(1-946)-3xFLAG his3-11,15 trp1-1 leu2-3,112 can1-100 bar1Δ MATa pep4::KANMX6* |
| AT27 | [a] W303 | *ade2-1 URA3::GAL-mrc1Δ(1-720)-3xFLAG his3-11,15 trp1-1 leu2-3,112 can1-100 bar1Δ MATa pep4::KANMX6 mrc1(1-720)::natMX6* |
| AT28 | [a] W303 | *ade2-1 URA3::GAL-mrc1Δ(1-946)-3xFLAG his3-11,15 trp1-1 leu2-3,112 can1-100 bar1Δ MATa pep4::KANMX6 mrc1(1-720)::natMX6* |
| AT29 | [a] W303 | *ade2-1 URA3::GAL-mrc1Δ(1-720)-3xFLAG-NLS his3-11,15 trp1-1 leu2-3,112 can1-100 bar1Δ MATa pep4::KANMX6 mrc1(1-946)::natMX6* |
| AT30 | [a] W303 | *ade2-1 URA3::GAL-mrc1Δ(1-946)-3xFLAG his3-11,15 trp1-1 leu2-3,112 can1-100 bar1Δ MATa pep4::KANMX6 mrc1(1-946)::natMX6* |
| AT37 | [a] W303 | *ade2-1 URA3::GAL-mrc1Δ(1-720)-3xFLAG-NLS his3-11,15 trp1-1 leu2-3,112 can1-100 bar1Δ MATa pep4::KANMX6 mrc1Δ::TRP1* |
| AT38 | [a] W303 | *ade2-1 URA3::GAL-mrc1Δ(1-946)-3xFLAG his3-11,15 trp1-1 leu2-3,112 can1-100 bar1Δ MATa pep4::KANMX6 mrc1Δ::TRP1* |
| JAY2087 | [b] CG379 | *MATαlpha MRC1* |
| AT13 | [b] CG379 | *MATαlpha MRC1::HphMX6* |
| AT14 | [b] CG379 | *MATαlpha mrc1(AQ)::HphMX6* |
| AT15 | [b] CG379 | *MATαlpha mrc1(1-720)::NatMX6* |
| AT16 | [b] CG379 | *MATαlpha mrc1Δ::NatMX6* |
| AT18 | [b] CG379 | *MATαlpha mrc1-K954E* |
| AT19 | [b] CG379 | *MATαlpha mrc1Δ(950-961)* |
| AT23 | [b] CG379 | *MATαlpha pol2-D1955K::NatMX6* |
| AT31 | [b] CG379 | *MATαlpha mrc1(1-946)::NatMX6* |

| Strain Name | Genetic background | Genotype |
|---|---|---|
| JAY3271 | [c] JAY291 | *MATa can1Δ::NatMX4 ura3 MRC1* |
| AT17 | [c] JAY291 | *MATa can1Δ::NatMX4 ura3 mrc1Δ::HphMX6* |

**Footnotes**

**a.** Haploid strains isogenic with the W303 background (Ralser et al. 2012) were used in all protein purifications and spot assays for HU sensitivity.

**b.** All haploid strains isogenic with the CG379 background shared the following genotype: *MATαlpha ade5-1*, *his7-2*, *leu2-3,112*, *Leu*$^{+}$, *ura3-52*, *trp1-289*, *cup1Δ*, *RSC30*, *sfa1Δ::hisG*, *PLM2::SFA1-V208I-CUP1-*$_{Kl}$*URA3-*$_{Sc}$*URA3-5'SFA1-BglII-KanMX4*. Specific *MRC1* mutations are noted in the table, and were all built by modifying the native *MRC1* locus on chromosome III. These strains were all derived from JAY2087 which is described in detail (McLaughlin et al. 2020).
They were used as haploids for measurements of the following spontaneous mutation rates, shown in the respective plots:
Fig. S9C - Interstitial deletion of the double *URA3* cassette inserted on the right arm of chromosome IV distal to *PLM2* (*PLM2::SFA1-V208I-CUP1-*$_{Kl}$*URA3-*$_{Sc}$*URA3-5'SFA1-BglII-KanMX4*; with counter selection for 5-FOA resistance.
Fig. S9D - Forward mutation to canavanine resistance at the native *CAN1* locus on the left arm of chromosome V; with counter selection for canavanine resistance.
Fig. S9E - Reversion of the *trp1-289* non-sense mutation allele, at native *TRP1* locus, centromere-linked on the right arm of chromosome IV; with selection for tryptophan prototrophy in Trp drop-out media.

**c.** The AT17 haploid strain (*MATa can1Δ::NatMX4 ura3 mrc1Δ::HphMX6*), which is isogenic with the JAY291 background (Argueso et al. 2009), was crossed to the various CG379 *MATαlpha* haploids. The resulting hybrid diploid strains were hemizygous at the *MRC1* locus (*i.e.*, the knockout allele from AT17 over the specific *MRC1* allele present in the respective CG379 haploid parent). The diploids were used for measurements of the following spontaneous LOH mutation rates, shown in the respective plots:
Fig. S9A - LOH at the double *URA3* cassette inserted on the right arm of chromosome IV distal to *PLM2*; with counter selection for 5-FOA resistance.
Fig. S9B - LOH at the *CAN1* locus on the left arm of chromosome V; with counter selection for canavanine resistance.

**Table S2: DNA oligonucleotides used in this study**

| Name | 5' to 3' Sequence |
|---|---|
| Mrc1 SQTQ Ala Donor A | GGAAATGGGCCCAATGACATAGACAATCCGCCAGAGTTGACTGGGA<br>ACGGGTTTTTATTTGCCAATGCCACATTAAATCGTGTTAAGAACAGAT<br>TAGAAGGCAAGAAAGCACCCGAACAGAATCACAATAATGGAAAAGAT<br>AGGAGTGAGAATTCGTTACCAGCTCAATTAATCTCTAATCTTTACGAT<br>GGTGGTGAAGAGCTGGAGAAATCCGAAGTTAAAGATAATAGTTACAG<br>TGAGAAAAATGTCTCTTCTTCATTCGCTCAGGCTCAAAGAATACCTGT<br>CAGCATCCAACAAGACAAGGTGTTTAACGTACCTATCCATTCCGTTAA<br>TGACGGGAAACCCGCTCAATTAATCAAAGAAGATGGTCTCGTCAACG<br>AAACTGCTCAAGCGCTGAAAACACCTCTCACGACAGGACGGCCAGG<br>GGCTGCTCAGCGTATAGATAGCAGCGGTGCAACCGCTCAAGCTCAG<br>CCTATAAAATCAATTGAACCACAAGCTCAAATAATTACGACTTCCAGC<br>AATCATTCTAATGCTCTGTCGCCGAAAATTCCTATAATTCCAACGGAA<br>CTTATTGGAACTAGTCCTTTATTTCAGTCCATTCAAAATCGTGGTCCT<br>GACGCTCAAATGGATGTACCACCACAGACTGCGCATGATGAGGATAA<br>GGCTCAAGCAATTGGTATTCCACAGGCTACGCATCAGGAACAGAAAG<br>CTCAAATAGATACAGTCGCACAAACACTTCAAGATGAAGTACCACACA<br>CATTAAAAATTCGGGAAATACAAAGCGAATTGGCTTCAGAAGATTCTA<br>AAAGGGAAAAAGCTAGAAATGTCGAATACAAAAAACCGCAAAAGCCA<br>ATACCAACCAAAAAATTTTTTTCTAAAGAGTCTTTTCTTGCTGATTTTG<br>ACGATTCGTCCTCAAATGAGGACGATGATATAAAGCTGGAAAATGCA<br>CATCCAAAACCAGTACAGAATGACGATGAACTGCATGAGAACAAAAG<br>TGTTGAATTAAATCTAACTGATGAGACAAGAATAAATGAGAAAAGGGT<br>TCCACTATTGAGCTCCTATGCTAATAATTTGAAACGGGAAATTGACTC<br>CTCGAAATGCATAACACTGGACCTCGACAGCGACAGTGACGAATATG<br>GAGATGATGATATGGATAGCATTAAATTATCCAAAGATGAAAGCGTTTT<br>GCCAATCGCTCAGTTATCAAAGGCGACTATATTAAACCTGAAGGCAA<br>GACTCTCTAAACAAAATCAAAAGTTGGCTCAAAGGCCGAATAAAAGTA<br>AAGACCCAAAAGTTGATCATAATGTACTATTAAATACCTTAAGGAAAGC<br>TAGCAGGAAACAAATTCTGGATCATCAGAAAGAAGTGATTGAAACTAA<br>GGGATTGAAATTGGAAGATATGGCCAAAGAAAAGGAAATAGTAGAGA<br>ACTTGCTAGAACAAGAAATTCTTAGAAATAAAAGAATCAGACAAAAAG<br>AGAAGAGAAGGG |
| Mrc1 SQTQ Ala Donor B | GAATCAGACAAAAAGAGAAGAGAAGGGAAAAACTTGAAGAAAATGAC<br>TTTCAATTGAACGCCCATGATTCCGGTTCTGACTCAGGGTCAGAGTC<br>TTCTGGATTTGCATTGAGTGGTAATGAAATTGCCGATTATGAATCATCT<br>GGTAGTGAAAATGACAATCGTAGAGAATCCGATAGTGAAAAGGAAGA<br>TGATGAAATTATTCTAAAGCAAAAAAAATCCCACCACGTGAAACATATC<br>ATTAACGAATCAGACTCTGATACCGAAGTCGAAGCCAAACCCAAAGA<br>GAAAGCTGATGAAAGTCTACCGAAAAGGATCGCCATTAATCTGGGCC<br>ATTACGGTGATAATATCGGAGAGGATACCGACAAATTTCAAGAAACCA<br>ATGTTCTAGATGCTCAAAACATTGAAGAGGTGATGGCCGAAAGAAATA<br>CAATCGAAAATGAAGTCAAAGATGATGTATACGTCAATGAAGAAGCTG<br>ATGAAGCAATTCGCCGTCAGCTGATTGACAAAGAGAAATTGCAACTA<br>AAACAGAAAGAAAAGGAGCACGAGGCTAAGATAAAAGAGTTGAAGA<br>AAAGAGGTGTCACAAACTTCTTTGAAATGGAAGCCGAAGAATCCGAG<br>GATGAATGGCATGGTATCGGTGGTGCAGATGGAGAAGGATCTGACG<br>ACTATGATTCTGACCTAGAAAAAATGATCGATGATTATTCCAAGAACAA<br>TTTCAATCCTCATGAAATCAGAGAAATGCTTGCCGCAGAAAATAAGGA |

| Name | 5' to 3' Sequence |
|---|---|
| | AATGGATATCAAAATGATAAACAAAATTCTTTATGACATTAAAAATGGAGGATTTCGTAATAAAAGAGCAAAAAATAGCTTAGAGCTAGAACTAAGTGATGATGATGAAGACGACGTACTACAACAATATCGTCTCAAAAGGAGAGAATTGATGAGAAAAAGGAGATTAGAAATTGGCGATGATGCAAAGCTTGTTAAAAACCCAAAGTCGAGCGCATTTTTTGAAAGTATGGTGGAAGATATTATCGAATATAAAAACCCCTTCGGGGCTGAAGAGGAATATAACCTAGACATTACAAGTACGGCTACTGACCTAGATGCTCAGGACAATAGCATTAATGTAGGTGATAATACGGGAAATAATGAACAGAAACCTGTAGATCAAAAGAACAAAAAGGTTATCATATCTGAAGACTTTGTGCAGAAAAGCTTATCCTTCCTAAAAAGTAATAATTATGAGGATTTTGAAACGGATAAGGAGCTATCAAGGATACAACATGGCAATGATGAAGCAATTGAAGATTTATATACCCTAAAACAGAATAGTTCGATAAAGTCATTCACAAATGCTCAAACTGATTCGACCACTTCAAAAACAGTAAACACAATAATTGACTTAGAAAAGCGCCCAGAGGATGAAGATGAGGTGGAAAATGGAGATACATCACTAGTTGGCG |
| Mrc1_K954E donor | GATAAGGAGCTATCAAGGATACAACATGGCAATGATGAAGCAATTGAAGATTTATATACCCTAGAACAGAATAGTTCGATAAAGTCATTCACAAATTCTCAAACTGATTCGACCACTTCAAAAACAGTAAACACAATAATAGATTTGGAGAAACGTCCTGAAGATGAAGATGAGGTGGAAAATGGAGATACATCACTAGTTGGCGTTTTCAAACATCCATCAATAATCAAATCCTTTGCGTCCAGAACAGACATAAATGATAAGTTTAAAGAAGGTAACAAGACAGTCAAGATTCTAAAGTCTTACAAGACAGTG |
| Mrc1_N16del donor | GATAAGGAGCTATCAAGGATACAACATGGCAATGATGAAGCAATTGAAGATTTCACAAATTCTCAAACTGATTCGACCACTTCAAAAACAGTAAACACAATAATAGATTTGGAGAAACGTCCTGAAGATGAAGATGAGGTGGAAAATGGAGATACATCACTAGTTGGCGTTTTCAAACATCCATCAATAATCAAATCCTTTGCGTCCAGAACAGACATAAATGATAAGTTTAAAGAAGGTAACAAGACAGTCAAGATTCTAAAGTCTTACAAGACAGTGGGGAGTTCAAAAGCTTCTATC |
| JAO270 (*CAN1* Forward) | GGTGTATGACTTATGAGGGTG |
| JAO273 (*CAN1* Reverse) | TATACTTATAGTTGGATCCAG |
